# Cryo-EM structure of CYP2C9 reveals a dimer-of-trimers assembly

**DOI:** 10.64898/2026.08.30.747968

**Authors:** Hiroki Tanino, Hirofumi Tsujino, Teruhiro Nakao, Chikara Oie, Fumiaki Makino, Tomoko Miyata, Kazuki Kasai, Keiichi Namba, Tsuyoshi Inoue

## Abstract

Human cytochrome P450 2C9 (CYP2C9) is a hepatic microsomal enzyme involved in the oxidative metabolism of clinically important drugs, but the structural organization of its oligomeric assemblies outside crystallographic packing environments remains poorly understood. Here, we report the cryo-EM structure of human CYP2C9 determined under aqueous, membrane-free conditions at 3.31 Å resolution. The structure reveals a C2-symmetric hexameric assembly organized as a dimer of trimers. Individual protomers retain the conserved P450 fold and heme-binding architecture observed in previously reported crystal structures, indicating that assembly formation does not substantially perturb the catalytic core. The hexamer is stabilized by defined intra-trimer interfaces involving the N-terminal region and residues around Trp212 and Phe482, together with inter-trimer interfaces involving Leu71 and the 220–227 loop. These interfaces are distinct from the crystal packing contacts observed in CYP2C9 crystal structures, demonstrating that the assembly is not a simple recapitulation of crystallographic packing. Notably, the inter-trimer interface is located near the FG-loop-containing surface previously implicated in membrane association. This suggests that the observed hexamer may represent a membrane-free association of two trimers through membrane-related surfaces, whereas the trimeric arrangement itself may be compatible with membrane-associated organization. The structure therefore provides a framework for investigating how trimer formation, membrane interaction and local conformational changes in the FG-loop region may influence CYP2C9 function.

## 1. Introduction

Cytochrome P450 enzymes constitute a large family of heme-containing monooxygenases that play diverse roles in the biosynthesis and metabolism of endogenous compounds, including steroid hormones, as well as in the metabolism of xenobiotics (Manikandan & Nagini, 2018; Guengerich, 2019). P450 enzymes catalyze substrate oxidation through heme-dependent activation of molecular oxygen, with the required reducing equivalents supplied by redox partner proteins. In eukaryotic microsomal P450 systems, the P450 catalytic domain is associated with the endoplasmic reticulum membrane and receives electrons from the membrane-bound NADPH-cytochrome P450 reductase (CPR) (Shukla *et al*., 2009; Barnaba *et al*., 2017). Among human drug-metabolizing P450 enzymes, CYP2C9 is an important hepatic isoform involved in the oxidative metabolism of clinically used drugs, including anticoagulants and non-steroidal anti-inflammatory drugs (Zanger & Schwab, 2013). Structural characterization of CYP2C9 is therefore essential for understanding its substrate recognition, ligand access pathways, and catalytic mechanism (Li & Poulos, 1996).

Crystal structures of human CYP2C9 have provided important insights into its protomer fold, heme-binding core, and active-site architecture, including substrate-free, S-warfarin-bound, and flurbiprofen-bound forms (Williams *et al*., 2003; Wester *et al*., 2004). These structures established the conserved P450 fold of CYP2C9 and revealed conformational features relevant to substrate recognition (Wester *et al*., 2004). However, because these structures were obtained using engineered soluble constructs and crystallographic packing environments, they do not directly describe how CYP2C9 is organized in solution or at the membrane interface. Previous biochemical and biophysical studies have shown that CYP2C9 can form homomeric and heteromeric P450–P450 complexes, and that such interactions can alter CYP2C9-mediated metabolism (Bostick *et al*., 2016). In particular, homomeric complex formation was reported to activate CYP2C9 metabolism, indicating that the oligomeric state of CYP2C9 can influence catalytic activity (Bostick *et al*., 2016). More broadly, P450–P450 interactions have been proposed to modulate drug metabolism through changes in CPR binding, substrate binding and/or substrate turnover (Reed & Backes, 2017). A putative CYP2C9 hexamer has also been proposed from the crystal lattice of the S-warfarin-bound structure, but whether CYP2C9 forms a defined solution-state hexamer and which interfaces stabilize such an assembly have remained unclear.

Because CYP2C9 functions as a membrane-associated enzyme, the molecular surfaces involved in oligomerization should ultimately be considered in relation to membrane interaction. However, it remains unclear whether CYP2C9 can form a defined higher-order assembly outside crystallographic packing environments and which molecular interfaces stabilize such an assembly.

Here, we report the cryo-EM structure of human CYP2C9 determined under aqueous conditions. The structure reveals a hexameric assembly organized as a dimer of trimers, providing direct structural visualization of a defined CYP2C9 higher-order assembly in solution. Structural comparison with a previously reported CYP2C9 crystal structure (PDB ID: 1OG2; Williams *et al*., 2003) reveals that the overall fold of individual protomers is largely preserved, whereas the cryo-EM assembly is stabilized by interfaces distinct from crystal packing contacts. Notably, the inter-trimer interface is located near a surface previously implicated in membrane interaction (Cojocaru *et al*., 2011; Berka *et al*., 2011), suggesting a possible relationship between CYP2C9 oligomerization and membrane-associated organization.

## 2. Methods

### 2.1. Protein expression and purification

Recombinant human CYP2C9 was prepared as previously reported (Miyamoto *et al*., 2015). The CYP2C9 gene lacking the N-terminal transmembrane region (residues 2–21) was cloned into the pET3a-based expression vector pBEX and transformed into Escherichia coli BL21 Gold (DE3) cells for protein expression and purification. The purity of the purified CYP2C9 was assessed by SDS–PAGE, which showed a single band consistent with previous reports. The concentration of CYP2C9 was determined by the pyridine hemochromogen assay based on the heme content. In addition, the UV–visible spectrum of the CO-bound form was measured, and the characteristic absorbance at 450 nm confirmed that the purified CYP2C9 was in its catalytically active form.

### 2.2. Mass photometry

The oligomeric state of purified CYP2C9 was assessed using a TwoMP mass photometer (Refeyn Ltd., Oxford, UK). The concentration of the CYP2C9 stock solution was determined by the pyridine hemochromogen assay. For each measurement, 19 µl of 100 mM potassium phosphate buffer, pH 7.4, was placed in the measurement well, followed by the addition of 1 µl of the CYP2C9 sample, yielding a final monomer-equivalent concentration of 122.5 nM. The monomer-equivalent concentration was calculated using a molecular mass of approximately 53.1 kDa for CYP2C9 containing one bound heme b cofactor. A buffer-only control was measured under the same conditions.

A 60-s movie was acquired using AcquireMP 2025 R1.2 (Refeyn Ltd.). Molecular-mass calibration was performed using thyroglobulin (669 kDa) and ovalbumin (44 kDa) from the Gel Filtration Calibration Kit HMW (28403842, Cytiva), together with bovine γ-globulin (158 kDa; G5009, Sigma-Aldrich). Landing events were detected and converted from interferometric contrast to molecular mass using DiscoverMP v2025 R1 (Refeyn Ltd.). The high-molecular-mass population was characterized by Gaussian fitting in DiscoverMP.

### 2.3. Cryo-EM grid preparation

Purified CYP2C9 was adjusted to 0.65 mg ml^−1^ in 100 mM potassium phosphate buffer pH 7.4 for cryo-EM grid preparation. For initial cryo-EM screening, 3 µl of the sample was applied to QUANTIFOIL R1.2/1.3 Cu 200 grids (Quantifoil Micro Tools GmbH) glow-discharged by a JEC-3000FC Auto Fine Coater (JEOL) at 7 Pa and 20 mA for 20 s before sample application. Grids were blotted using a Vitrobot Mark IV device (Thermo Fisher Scientific) at 8°C and 100% humidity, with a blotting time of 2.5 s and a blot force of 0, and were then plunge-frozen in liquid ethane.

For the final dataset, 3 µl of the CYP2C9 sample was applied to EG-grid® without glow discharge. Grids were blotted using Vitrobot at 8°C and 100% humidity, with a blotting time of 1.0 s and a blot force of −15, and were plunge-frozen in liquid ethane.

### 2.4. Cryo-EM data collection

Cryo-EM data were collected using a JEM-Z200FSC transmission electron microscope (JEOL) operated at 200 kV and equipped with a K3 direct electron detector (Gatan). Movies were recorded in counting mode at a pixel size of 0.83 Å per pixel and fractionated into 40 frames. Automated data collection was controlled using SerialEM (Mastronarde, 2005), and carbon holes were detected using yoneoLocr (Yonekura *et al*., 2021).

For QUANTIFOIL R1.2/1.3 Cu 200 grids, two datasets were collected. The first dataset comprised 6806 movies recorded with an exposure time of 3.980 s, and the second dataset comprised 3389 movies recorded with an exposure time of 3.828 s. Both Quantifoil datasets were collected with a nominal defocus range of −0.7 to −2.2 µm and a total electron exposure of 80 e^−^ Å^−2^. For the EG-grid dataset used for final reconstruction, 17065 movies were recorded with an exposure time of 1.762 s, a nominal defocus range of −0.7 to −1.7 µm, and a total electron exposure of 40 e^−^ Å^−2^.

### 2.5. Image processing and three-dimensional reconstruction

All cryo-EM data were processed in CryoSPARC v4.7.0 (Punjani *et al*., 2017). Movies were corrected for beam-induced motion using Patch Motion Correction, and CTF parameters were estimated using Patch CTF Estimation.

For the two Quantifoil datasets, exposures were curated independently, followed by blob-based particle picking, particle extraction and iterative 2D classification. Selected particles from the two datasets were combined and subjected to ab initio reconstruction, heterogeneous refinement and non-uniform refinement. Although a hexamer-like reconstruction was obtained, the reconstruction remained strongly affected by preferred particle orientation and was not used for final structure determination (Supplementary Fig. S1(a)).

For the EG-grid dataset, exposures were curated following motion correction and CTF estimation. An initial subset of exposures was subjected to blob picking, particle extraction and iterative 2D classification. The selected particles were used for ab initio reconstruction, heterogeneous refinement and non-uniform refinement to obtain an initial oligomeric reconstruction, from which 2D templates were generated using Create Templates for template-based particle picking. The resulting particles were further cleaned by iterative 2D classification, ab initio reconstruction and heterogeneous refinement. Selected particles were subsequently subjected to reference-based motion correction, global and local CTF refinement and non-uniform refinement. The final reconstruction was obtained from 315,418 particles with C2 symmetry (Supplementary Fig. S2). The map resolution was estimated to be 3.31 Å using the gold-standard FSC = 0.143 criterion. The final map was sharpened in CryoSPARC using an automatically estimated sharpening B factor of 144.2 Å².

### 2.6. Model building and refinement

An initial model was prepared from chain A of the CYP2C9 crystal structure PDB ID 1OG2 and fitted into the cryo-EM density map using UCSF ChimeraX (Meng *et al*., 2023; Pettersen *et al*., 2021). The model was manually adjusted in Coot and refined against the cryo-EM map using phenix.real_space_refine (Afonine *et al*., 2018; Liebschner *et al*., 2019). The heme prosthetic group was modeled as heme b (HEM), with ligand restraints generated in Coot (Emsley *et al*., 2010). Manual model adjustment and real-space refinement were performed iteratively. The final model was validated by MolProbity (Chen *et al*., 2010) in Phenix, and refinement and validation statistics are summarized in Table 1. Engineered substitutions present in 1OG2 were reverted to the corresponding residues in the CYP2C9 construct used in this study before refinement.

**Table 1.** Cryo-EM data collection, refinement and validation statistics for the CYP2C9 hexamer structure.

| **Data collection and processing** |  |
| --- | --- |
| Microscope | JEM-Z200FSC |
| Voltage(kV) | 200 |
| Detector | K3 |
| Pixel size (Å) | 0.83 |
| Electron exposure (e<sup>-</sup>/Å<sup>-2</sup>) | 40 |
| Symmetry imposed | C2 |
| Final particle images (no.) | 315,418 |
| Map resolution (Å) | 3.31 |
| FSC threshold | 0.143 |
| Map sharpening B factor (Å<sup>2</sup>) | 144.2 |
| **Refinement** |  |
| Initial model used | PDB: 1OG2, modified to match the CYP2C9 sequence used in this study |
| Model resolution (Å) | 3.6 |
| FSC threshold | 0.5 |
| **Model composition** |  |
| Non-hydrogen atoms | 22,388 |
| Protein residues | 2,766 |
| Ligands | 6 HEM |
| **B factors (Å<sup>2</sup>)** |  |
| Protein | 178.25 |
| Ligand | 153.86 |
| **R.m.s. deviations** |  |
| Bond lengths (Å) | 0.004 |
| Bond angles (°) | 0.952 |
| **Validation** |  |
| MolProbity score | 2.52 |
| Clashscore | 43.08 |
| Ramachandran plot |  |
| Favored (%) | 93.97 |
| Allowed (%) | 5.70 |
| Outliers (%) | 0.33 |
| Rotamer outliers (%) | 0.96 |
| Cβ outliers (%) | 0.00 |
| **Map–model correlation** |  |
| CC(mask) | 0.86 |
| CC(volume) | 0.85 |
| CC(peaks) | 0.79 |

### 2.7. Map and model validation

Directional resolution anisotropy of the final cryo-EM reconstruction was evaluated using the Remote 3DFSC Processing Server (Tan *et al*., 2017). The two independently refined half maps were used to calculate directional Fourier shell correlations, and the resulting directional FSC distribution and sphericity were used to assess anisotropy of the reconstruction.

Local map–model agreement was evaluated using the DAQ-score Free Server provided by the Kihara laboratory (Terashi *et al*., 2022). The final CYP2C9 atomic model and cryo-EM density map were submitted to the server to calculate residue-wise DAQ scores. For visualization of local variations in model quality, residues with DAQ scores below 0.2 were highlighted, and the distribution of these residues was compared among individual subunits.

## 3. Results

### 3.1. Overall structure of the CYP2C9 dimer-of-trimers assembly

To assess whether purified CYP2C9 forms oligomeric species in solution, mass photometry was performed at a final CYP2C9 concentration of 122.5 nM, calculated on a monomer-equivalent basis. CYP2C9 has a theoretical monomeric molecular mass of 52.4 kDa, or approximately 53.1 kDa when one bound heme *b* cofactor is included. The mass distribution contained a prominent low-molecular-mass population in the approximate monomeric mass range, together with a distinct high-molecular-mass population centered at approximately 354 kDa (Fig. 1a). No comparable high-molecular-mass signal was detected in the buffer-only control. Although the measured mass did not permit assignment of a unique oligomeric stoichiometry, the detection of this population supports the presence of higher-order CYP2C9 oligomers in solution. Accordingly, the mass photometry data were interpreted as qualitative evidence for oligomerization rather than as a definitive determination of the complete oligomeric-state distribution.

**Figure 1.**
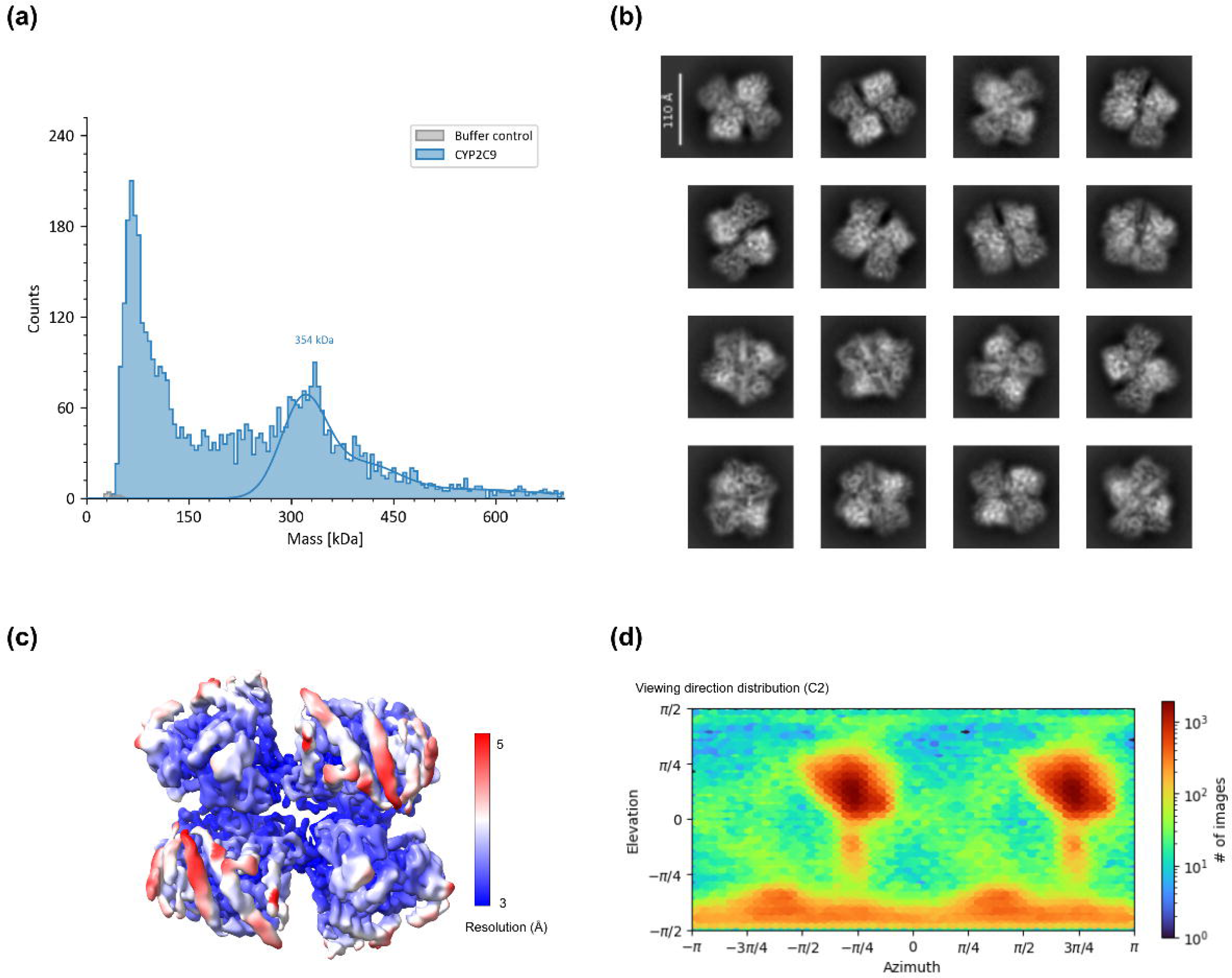
Solution-state oligomerization and cryo-EM analysis of CYP2C9. (a) Mass distribution of purified CYP2C9 measured at a final monomer-equivalent concentration of 122.5 nM (blue), together with a buffer-only control (gray). A prominent low-molecular-mass population and a distinct high-molecular-mass population centered at approximately 354 kDa were detected. The solid blue curve represents a Gaussian fit to the high-molecular-mass population. No comparable high-molecular-mass population was detected in the buffer-only control. (b) Representative 2D class averages of CYP2C9 particles collected on an EG-grid. The class averages show triangular and ring-like features consistent with oligomeric particles. Scale bar, 110 Å. (c) Cryo-EM map of the CYP2C9 assembly colored by local resolution. (d) Viewing-direction distribution of the particles contributing to the final *C*2 reconstruction.

To structurally characterize these oligomeric particles, cryo-EM analysis was initially performed using a conventional Quantifoil Cu R1.2/1.3 grid. Two-dimensional classification revealed particle images substantially larger than expected for monomeric CYP2C9, which measures approximately 60–70 Å along its longest axis. The resulting class averages included triangular and ring-like features with apparent symmetry, consistent with oligomeric assemblies (Supplementary Fig. S1(b)). These observations were also consistent with the mass photometry data indicating the presence of higher-order CYP2C9 species. However, pronounced preferred particle orientation limited the three-dimensional reconstruction to low resolution and prevented reliable high-resolution structural interpretation (Supplementary Fig. S1(a)). To improve the particle orientation distribution, data were subsequently collected using an epoxidized graphene grid (EG-grid), which provides a functionalized graphene support that adsorbs protein particles and has been reported to improve particle density and orientation distribution (Fujita *et al*., 2023).

The use of the EG-grid improved the particle orientation distribution and enabled successful three-dimensional reconstruction. The final *C*2-symmetric reconstruction was obtained from 315,418 particles at an overall resolution of 3.31 Å, based on the gold-standard FSC = 0.143 criterion. The map showed well-resolved secondary-structure features and broad viewing-direction coverage (Fig. 1c,d). Directional FSC analysis using 3DFSC yielded a sphericity value of 0.967, indicating that the reconstruction did not exhibit severe directional anisotropy (Supplementary Fig. S3).

Orthogonal and rotated views of the cryo-EM structure of CYP2C9 (CYP2C9^EM^) illustrate the relative orientation of the two trimers and the overall symmetry of the assembly (Fig. 2a,b). In particular, the side view clearly shows that the hexamer is organized as a dimer of trimers (Fig. 2b). For clarity, the three protomers in one trimer are designated A, B and C, whereas the corresponding protomers in the opposing trimer are designated A′, B′ and C′, corresponding to PDB chain IDs D, E and F, respectively.

**Figure 2.**
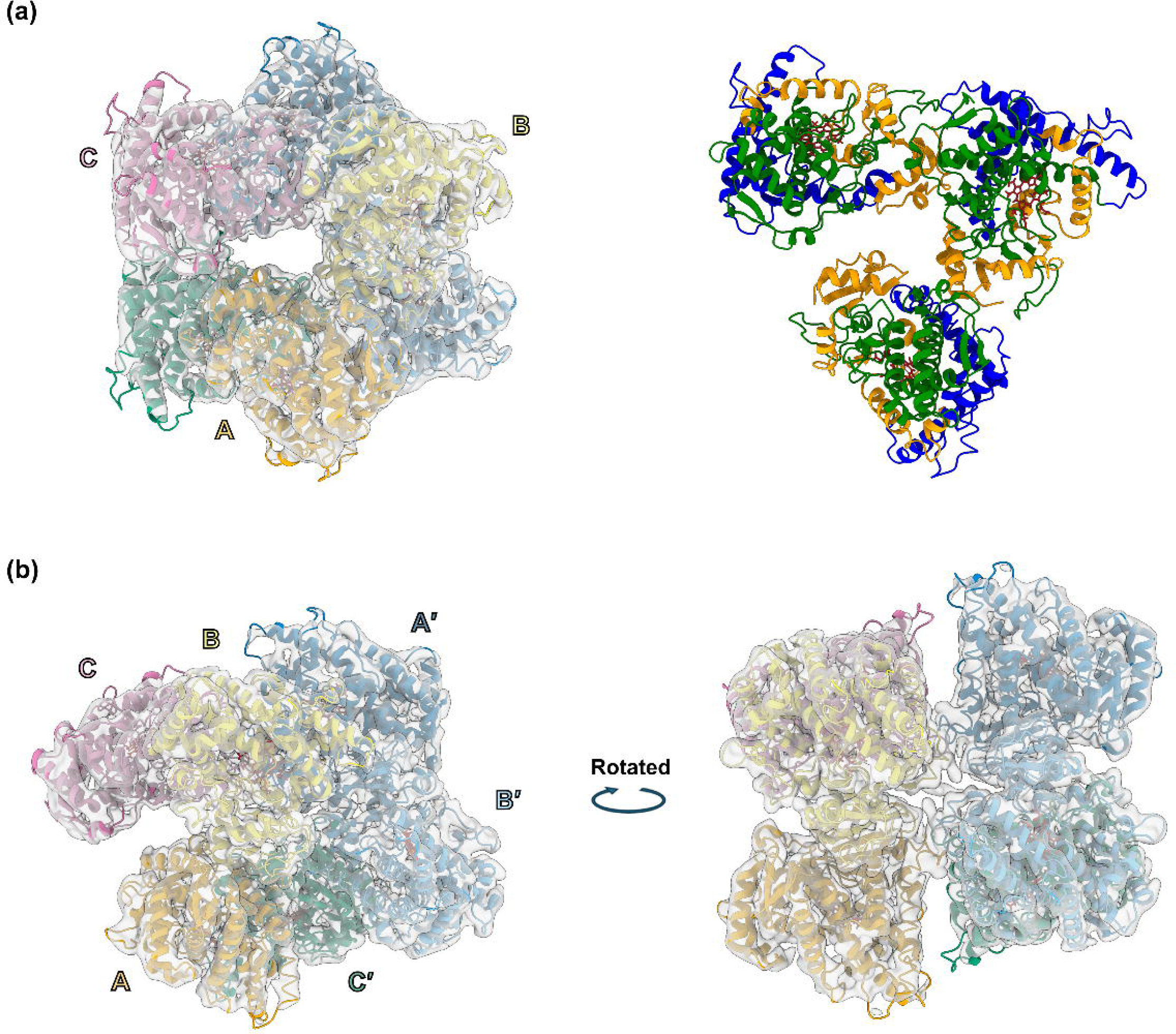
Overall architecture of the CYP2C9 hexamer. (a) Cryo-EM map and atomic model of the CYP2C9 hexamer. Individual subunits are shown in different colors, and heme groups are shown as red sticks. The right panel shows a domain-colored representation of a single CYP2C9 trimer extracted from the hexamer. The N-terminal/β-rich region, substrate-recognition region and heme-binding core are colored orange, blue and green, respectively. (b) Hexameric assembly of CYP2C9 shown in two orientations. The structure consists of two opposing trimers arranged in a face-to-face manner. Subunits are labeled A–C and A′–C′, corresponding to the two trimeric units.

Within each trimer, three subunits form an asymmetric triangular arrangement around a pseudo-threefold axis (Fig. 2a). The A–B and B–C interfaces are relatively tight, whereas the A–C interface is more open, indicating that the trimer does not possess strict *C*3 symmetry. This asymmetric contact pattern likely contributes to the pseudo-threefold organization of the trimeric unit. Each CYP2C9 monomer retains the canonical cytochrome P450 fold, comprising the N-terminal/β-rich region, substrate-recognition region, and the heme-binding core (Fig. 2a). These structural features are preserved in the oligomeric assembly, indicating that multimerization does not substantially perturb the overall fold of individual subunits.

Local model quality was further evaluated using DAQ-score analysis, which supported the overall consistency of the atomic model with the cryo-EM density, while revealing local variation in model-to-map agreement (Supplementary Fig. S4).

### 3.2. Comparison with previously reported CYP2C9 crystal structures

To assess in detail whether oligomerization affects the structure of CYP2C9, individual subunits in the cryo-EM model (chains A, B, and C) were compared with a previously reported crystal structure (PDB ID: 1OG2) (Williams *et al*., 2003). The crystal structure contains two CYP2C9 molecules in the asymmetric unit. Since chains A and B of 1OG2 are nearly identical, with a Cα root-mean-square deviation (RMSD) of 0.356 Å over 461 aligned residues, chain A was used as the representative chain for structural comparisons. Structural superposition of the cryo-EM subunits with the crystal structure revealed a high degree of overall similarity, with Cα RMSD values of 1.36 Å for chain A, 0.86 Å for chain B and 1.24 Å for chain C, each calculated over 461 aligned residues (Fig. 3a; Supplementary Fig. S6). The overall architecture of CYP2C9 is therefore largely preserved in the cryo-EM structure, including the characteristic cytochrome P450 fold and the heme-binding core. Despite minor local differences in surface-exposed loop regions, including residues around Ser210–Asn231 in some subunits (Supplementary Fig. S6), the position and orientation of the heme prosthetic group are well conserved between the cryo-EM and crystal structures (Fig. 3b). These results indicate that hexamer formation does not substantially alter the tertiary structure of individual CYP2C9 subunits, although local conformational differences are observed in surface-exposed regions involved in oligomerization interfaces, particularly around the membrane-associated FG-loop region described below (Fig. 5).

**Figure 3.**
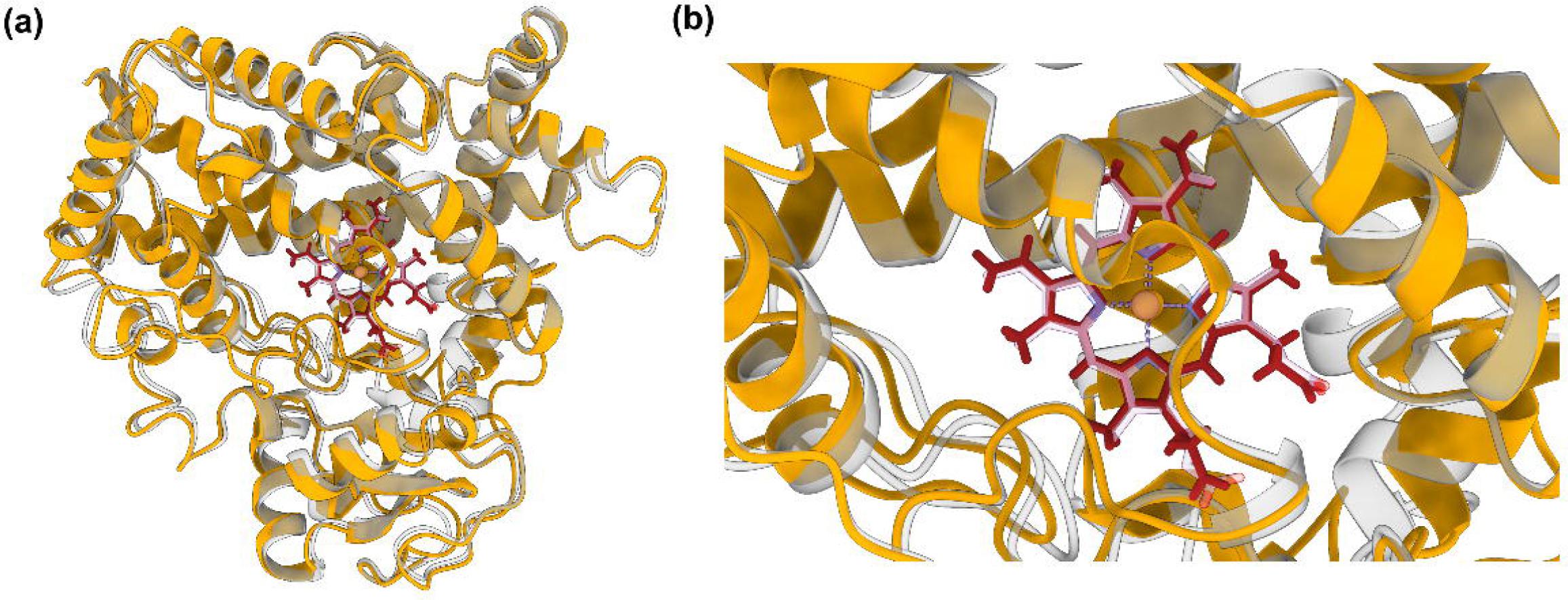
Comparison of the cryo-EM structure with the crystal structure of CYP2C9. (a) Structural superposition of chain A from the CYP2C9 cryo-EM model and chain A of the CYP2C9 crystal structure (PDB ID: 1OG2). The cryo-EM model and crystal structure are shown in orange and gray, respectively, highlighting their overall structural similarity. Heme is shown as red sticks. (b) Close-up view of the heme-binding site. The position and orientation of the heme prosthetic group and the surrounding structural elements are largely conserved between the cryo-EM and crystal structures. Heme is shown as red sticks and the iron atom as a sphere.

### 3.3. Structural basis of intra- and inter-trimer interactions

The hexameric assembly of CYP2C9 is organized as a dimer of trimers, in which both intra- and inter-trimer interfaces contribute to the overall architecture (Fig. 4a).

**Figure 4.**
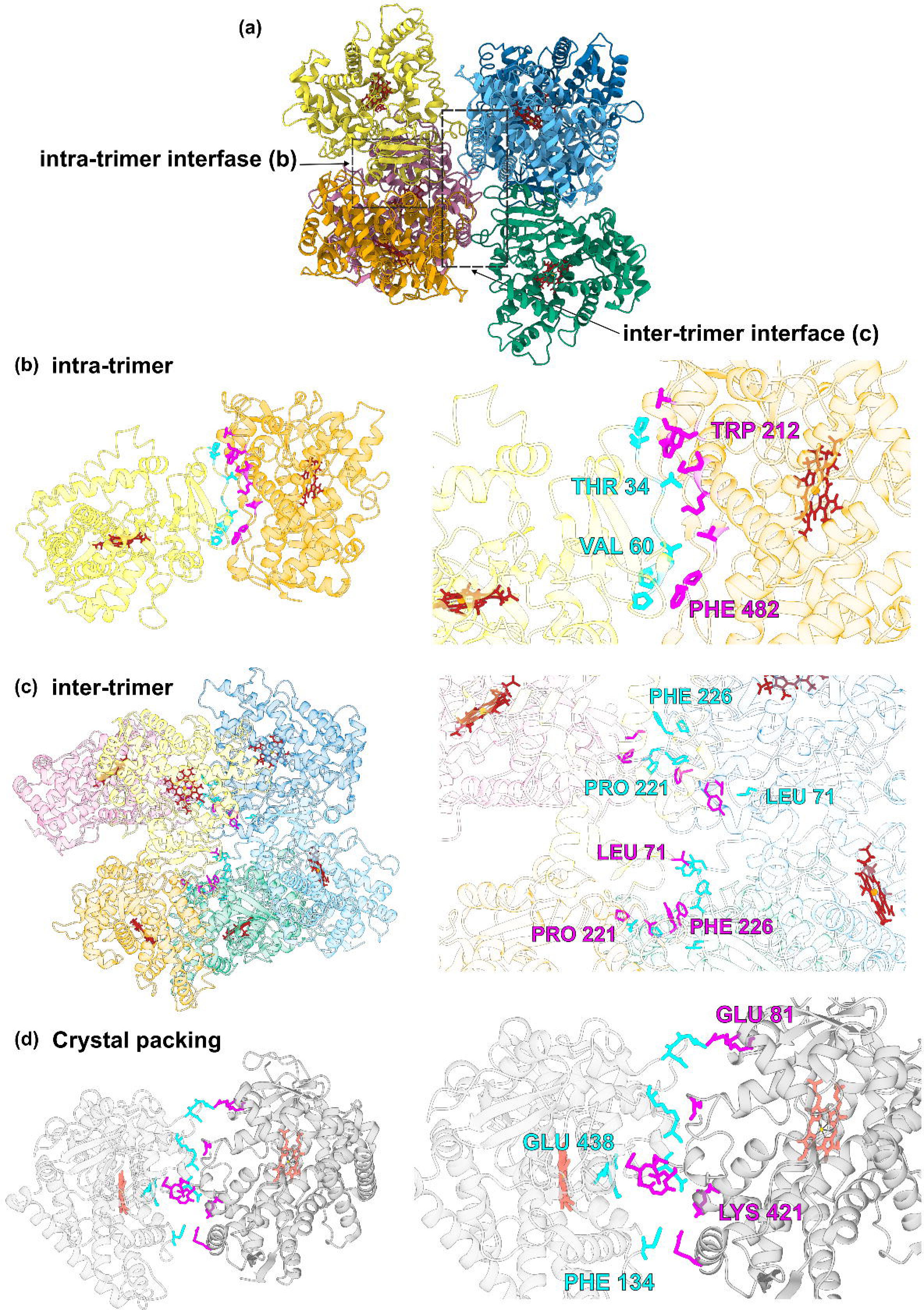
Intra- and inter-trimer interfaces of the CYP2C9 hexamer and comparison with crystal packing contacts. (a) Overall architecture of the CYP2C9 hexamer, showing the positions of the intra-trimer and inter-trimer interfaces. The regions enlarged in (b) and (c) are indicated. Each protomer is shown in a different color, and heme groups are shown as red sticks. (b) Intra-trimer interface between adjacent protomers (between chains A and B) within a single trimer. The left panel shows an overview of the contacting protomers, and the right panel shows a close-up view of the interface. Interface residues from the two protomers are respectively shown as magenta (in chain A) and cyan sticks (in chain B), with representative residues labeled. (c) Inter-trimer interface between the two opposing trimers. The left panel shows an overview of the inter-trimer interface, with interface residues highlighted as sticks. The right panel shows a close-up view of representative inter-trimer contacts. Interface residues from the two opposing trimers are respectively shown in magenta (in trimer ABC) and cyan (in trimer A′B′C′), and representative residues including Leu71, Pro221 and Phe226 are labeled. (d) Crystal packing interface observed in the CYP2C9 crystal structure (PDB ID: 1OG2). Two adjacent molecules in the asymmetric unit are shown in gray, with interface residues highlighted as sticks in magenta and cyan. Representative residues Glu81, Lys421, Phe134 and Glu438 are labeled. The residues forming the cryo-EM intra- and inter-trimer interfaces are distinct from those involved in this crystal packing interface.

Within each trimer, adjacent protomers form a well-defined intra-trimer interface (Fig. 4b). This interface is formed by an extended contact surface between the 206–212 region and the C-terminal segment of one protomer and the N-terminal region, including residues around 32–37 and 59–63, of the adjacent protomer. Representative interface residues include Trp212 and Phe482 from one protomer and Thr34 and Val60 from the adjacent protomer. The intra-trimer interfaces were observed between adjacent protomers, whereas no significant interface was detected between non-adjacent protomers such as chains A and C, indicating an open trimeric arrangement rather than a closed threefold-symmetric trimer.

In addition to the intra-trimer contacts, a distinct inter-trimer interface is formed between the two opposing trimers (Fig. 4c). This interface is mainly formed by Leu71 and residues in the 220–227 loop, including Pro221, Asp224, Tyr225, Phe226 and Pro227. These residues were detected at multiple contacts between the two opposing trimers, suggesting that this surface contributes to formation of the hexameric assembly. Pairwise inspection of the inter-trimer contacts from the ABC trimer further revealed an asymmetric contact pattern: chains A and B each contact more than one subunit in the opposing trimer, whereas chain C contacts only chain A′ (Supplementary Fig. S7).

The inter-trimer interface is located near the FG-loop region (residues 208–230), which has previously been implicated in CYP2C9–membrane interactions (Cojocaru *et al*., 2011; Supplementary Fig. S8). Thus, the inter-trimer interface partially overlaps with a membrane-related surface of CYP2C9.

To assess whether the interfaces observed in the cryo-EM structure correspond to crystal packing contacts, we compared them with the intermolecular contact observed in the CYP2C9 crystal structure (PDB ID: 1OG2). In this crystal structure, the packing contact involves residues such as Glu81 and Lys421 from one molecule and Phe134 and Glu438 from the adjacent molecule (Fig. 4d). These residues are located in regions distinct from those mediating both the intra- and inter-trimer interfaces in the cryo-EM assembly. Importantly, residues that contribute to the interfaces in the cryo-EM structure, including Trp212, Val60, Phe482, Leu71 and residues in the 220–227 loop, are not the principal residues involved in the crystal packing contact. This indicates that the interfaces observed in the cryo-EM structure represent a distinct mode of association rather than simply recapitulating crystal packing interactions.

Although the overall CYP2C9 fold is largely conserved relative to the crystal structure, local conformational differences were observed in the surface-exposed region around residues Ser210–Asn231. Notably, this region contributes to both intra- and inter-trimer contacts in the cryo-EM assembly, including residues around Trp212 and the 220–227 loop. These observations suggest that local rearrangements in this surface region may accompany oligomer formation in solution. A putative CYP2C9 hexamer was previously proposed from the crystal lattice of the S-warfarin-bound CYP2C9 structure (PDB ID: 1OG5), in which three side-to-proximal dimers assemble with approximate threefold symmetry to form a trimer-of-dimers architecture (Reed & Backes, 2017; Supplementary Fig. S9). This arrangement is fundamentally different from the dimer-of-trimers architecture observed in the present cryo-EM structure. Moreover, the crystal-lattice-derived model is based on side-to-proximal and side/distal contacts rather than the intra-trimer and FG-loop-associated inter-trimer interfaces identified in the cryo-EM assembly. Thus, although both structures suggest an intrinsic propensity of CYP2C9 to form higher-order assemblies, the present hexamer represents a distinct mode of oligomerization from the previously proposed crystal-lattice-derived model.

### 3.4. Membrane-related features of the oligomerization interface

The inter-trimer interface involves the 220–227 loop, which lies within the FG-loop region of CYP2C9 (Fig. 5a). Previous molecular dynamics simulations of membrane-bound CYP2C9 showed that the FG-loop region contributes to membrane interaction and influences the orientation and dynamics of CYP2C9 on the lipid bilayer (Cojocaru *et al*., 2011). Therefore, the inter-trimer interface observed in the CYP2C9^EM^ structure is thought to correspond to a surface that interacts with membrane. The hydrophobic properties of the membrane-binding surface may promote intermolecular association when this region is exposed to the aqueous environment, thereby stabilizing the inter-trimer contacts observed in the CYP2C9^EM^ structure.

**Figure 5.**
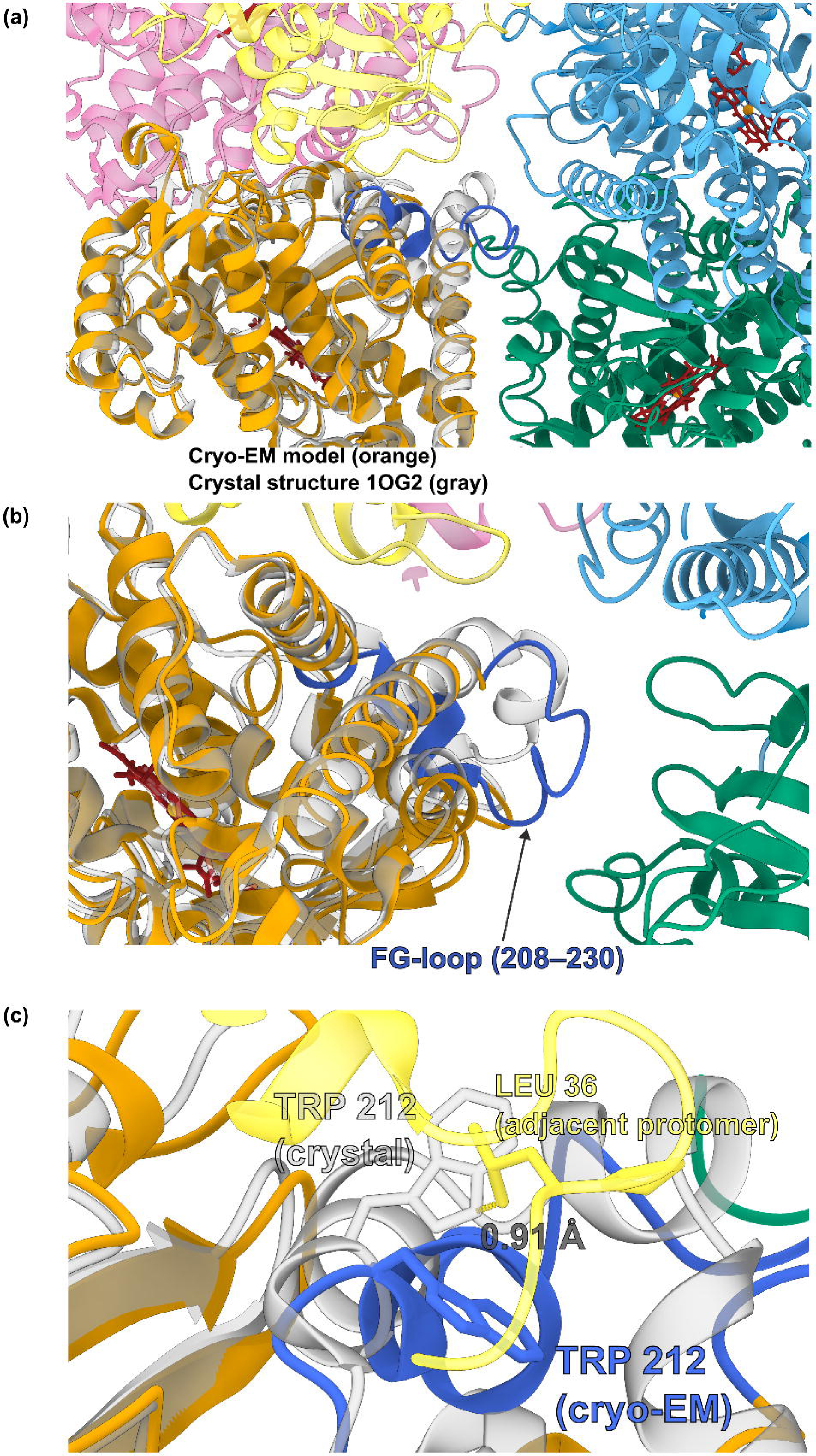
Conformational rearrangement of the FG-loop region at the CYP2C9 oligomerization interface. (a) Location of the FG-loop region (residues 208–230) in the CYP2C9 hexamer. Chain A is shown in orange, with its FG-loop region highlighted in royal blue. Other protomers are shown in distinct colors, and heme groups are shown as red sticks. The FG-loop region is positioned near the inter-trimer interface of the hexameric assembly. (b) Superposition of chain A from the cryo-EM structure with the previously reported CYP2C9 crystal structure. The cryo-EM model is shown in orange, with the FG-loop region highlighted in royal blue, and the crystal structure is shown in gray. The FG-loop region adopts a locally shifted conformation in the cryo-EM assembly relative to the crystal structure. (c) Close-up view of the Trp212 region. The position of Trp212 observed in the crystal structure is shown in gray, whereas that observed in the cryo-EM structure is shown in royal blue. Leu36 from the adjacent protomer is shown in yellow. The minimum interatomic distance between the crystal-state Trp212 and Leu36 of the adjacent protomer after superposition is 0.91 Å.

Structural comparison with the CYP2C9 crystal structure showed that this region adopts a locally shifted conformation in the cryo-EM assembly. The region showing local deviation from the crystal structure, particularly around residues 211–230, overlaps with the FG-loop/220–227 region involved in the oligomerization interface (Fig. 5b). Thus, the local structural difference observed in this surface-exposed region is spatially linked to the interface-forming surface of the hexamer.

A close-up comparison around Trp212 suggests that this local rearrangement may be important for accommodating the observed trimer interface. When the crystal structure was superposed onto the cryo-EM trimer, the position of Trp212 in the crystal structure showed overlap with Leu36 of the adjacent protomer, with a nearest atom distance of 0.91 Å (Fig. 5c). By contrast, in the cryo-EM structure, Trp212 is repositioned within the FG-loop region, avoiding this steric hindrance and allowing the adjacent protomers to form the observed intra-trimer interface. These observations suggest that the FG-loop region undergoes local conformational rearrangement associated with oligomerization.

Together, these findings identify the FG-loop region as a structural link between membrane-associated surfaces of CYP2C9 and the oligomerization interfaces observed in the cryo-EM structure. The overlap between the locally shifted FG-loop region, the Trp212-containing intra-trimer interface and the membrane-related 220–227 region suggests that this surface is structurally adaptable and may participate in both membrane interaction and protein–protein association.

## 4. Discussion

### 4.1. Overall structural organization of CYP2C9 assemblies

In this study, we determined the cryo-EM structure of human CYP2C9 and identified a hexameric assembly organized as a dimer of trimers. To our knowledge, this structure provides the first direct structural visualization of a CYP2C9 hexameric assembly in solution. While previous structural studies have primarily described individual CYP2C9 protomers in crystallographic environments, the present structure reveals a defined higher-order organization under aqueous, membrane-free conditions. Because the interface between the two trimers involves surfaces associated with membrane interaction, the complete hexamer may represent a membrane-free association of two trimeric units, whereas the trimeric arrangement itself may constitute a membrane-compatible organizational unit of CYP2C9.

Despite oligomer formation, the overall CYP2C9 fold remains highly similar to that observed in the crystal structure, with low Cα RMSD values for all three cryo-EM subunits. Structural differences are instead concentrated in surface-exposed regions involved in intermolecular interactions, particularly around the FG-loop region. Thus, the present structure suggests that oligomerization is associated primarily with local structural adaptation at interaction surfaces rather than with substantial rearrangement of the CYP2C9 catalytic core.

### 4.2. Distinct interfaces from crystal packing contacts

A putative CYP2C9 hexamer was previously proposed from the crystal lattice of the S-warfarin-bound structure (PDB ID: 1OG5), in which three side-to-proximal dimers associate with approximate threefold symmetry to form a trimer-of-dimers architecture (Reed & Backes, 2017; Supplementary Fig. S9). By contrast, the present cryo-EM structure is organized as a dimer of asymmetric trimers and is stabilized by a distinct set of intra- and inter-trimer contacts. The intermolecular interfaces identified in the cryo-EM assembly therefore differ not only in their constituent residues but also in their overall mode of oligomeric organization from the previously proposed crystal-lattice-derived model.

### 4.3. Solution-state assembly and membrane-related interfaces

The present structure was determined under aqueous, membrane-free conditions and therefore does not directly represent the native membrane-associated state of CYP2C9. A notable feature of the hexamer is that the inter-trimer interface involves the FG-loop region and adjacent surface elements previously implicated in CYP2C9–membrane interaction (Cojocaru *et al*., 2011). This region also undergoes local conformational rearrangement relative to the crystal structure. In particular, the Trp212-containing region adopts a conformation compatible with the observed intra-trimer interface, whereas superposition of the crystal-state conformation onto the cryo-EM trimer would generate severe steric overlap with the adjacent protomer. These observations suggest that the FG-loop region represents a conformationally adaptable surface capable of participating in both membrane interaction and protein–protein association. However, because the crystal construct used for comparison contains engineered substitutions within this region, the observed local conformational difference cannot be attributed exclusively to oligomerization.

This interpretation suggests that the hexameric assembly observed here may represent a membrane-free association rather than a direct representation of the physiological oligomeric state of CYP2C9. In a membrane environment, the FG-loop-containing surface would be expected to interact with or lie near the lipid bilayer. In the absence of a membrane, this surface may instead participate in protein–protein interactions, thereby promoting association between two trimeric units to form the observed hexamer.

Importantly, conformational changes in the FG-loop region may also have functional consequences for substrate access and catalytic activity. Previous molecular dynamics simulations showed that the opening and closing of substrate-access and product-release tunnels in CYP2C9 are influenced by the conformation of the FG loop, ligand binding and membrane association (Cojocaru *et al*., 2011). In particular, a membrane-directed substrate-access tunnel was more open when the FG loop adopted an extended conformation, and residues Ser220 and Pro221 within the FG-loop region contribute to the entrance of this tunnel. These observations suggest that the oligomerization-associated rearrangement of the FG-loop region observed in the present cryo-EM structure could alter substrate access to the buried active site and thereby influence CYP2C9 catalytic activity. Although the present structure does not directly establish such a functional effect, it raises the possibility that oligomerization and membrane association may modulate CYP2C9 activity through conformational changes in the FG-loop region.

## 5. Conclusion

Our results suggest that the trimeric arrangement, rather than the complete hexameric assembly, may represent a structurally meaningful organizational unit of CYP2C9. The hexamer observed under the present membrane-free aqueous conditions can be interpreted as an association of two trimers through surfaces that would otherwise be available for interaction with the lipid bilayer. In contrast, the intra-trimer contacts can be maintained without burying this membrane-facing surface, raising the possibility that the trimeric arrangement is compatible with membrane association. Furthermore, the involvement and local rearrangement of the FG-loop region suggest a potential structural link between trimer formation, membrane interaction and regulation of substrate access. Although the physiological relevance of the trimeric arrangement remains to be established, the present structure provides a framework for investigating how CYP2C9 molecules are organized on the endoplasmic reticulum membrane and how such organization may influence enzymatic function. Future studies using membrane-mimetic systems and interacting partners such as CPR will be required to determine whether the trimeric arrangement is retained in a membrane environment and how it affects CYP2C9 activity.

## Acknowledgements

This work was performed in part under the Collaborative Research Program of the Institute for Protein Research, Osaka University, CR-25-02.

## Data availability

The cryo-EM density map and atomic coordinates have been deposited in the Electron Microscopy Data Bank and Protein Data Bank, respectively. The raw micrographs collected in this study and used for structural analysis have been deposited in the Electron Microscopy Public Image Archive. Accession codes will be provided upon release.

