## Supporting Information for "Cryo-EM structure of CYP2C9 reveals a dimer-of-trimers assembly"

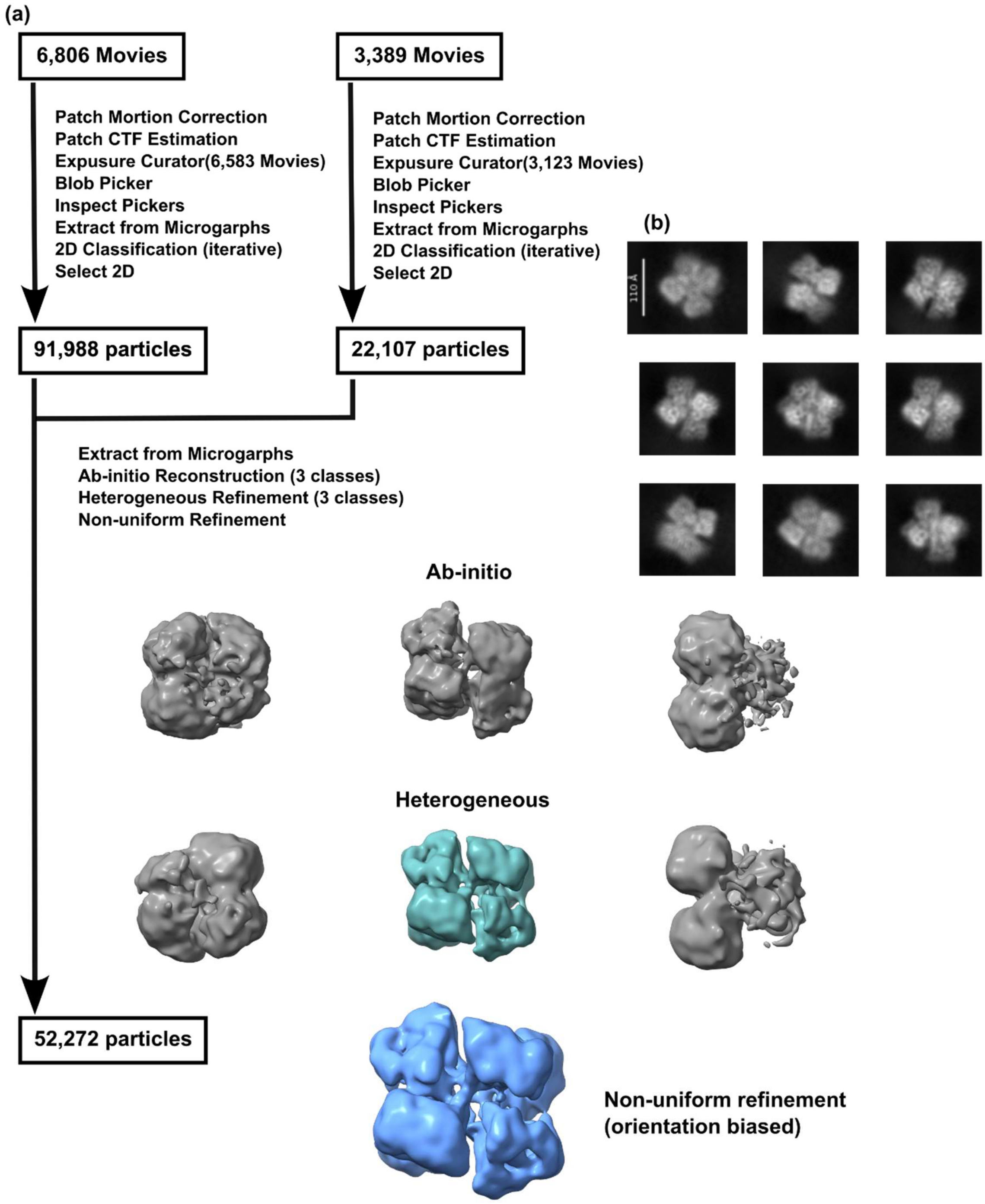


**Supplementary Figure S1. Cryo-EM analysis of CYP2C9 particles on conventional Quantifoil grids.**

(a) Processing workflow for the two Quantifoil datasets. A total of 6806 and 3389 movies were collected in the two independent datasets, respectively. After patch motion correction, CTF estimation, exposure curation, particle picking and iterative 2D classification, 91,988 and 22,107 particles were selected from the two datasets and combined for downstream 3D analysis. Ab initio reconstruction and heterogeneous refinement yielded oligomeric maps consistent with higher-order CYP2C9 assemblies. Subsequent non-uniform refinement produced a hexamer-like map; however, the reconstruction remained strongly affected by preferred particle orientation and was therefore not used for final structure determination. (b) Representative 2D class averages from the Quantifoil datasets. The class averages show triangular and ring-like particle images substantially larger than monomeric CYP2C9, consistent with oligomeric particles. Scale bar, 110 Å.

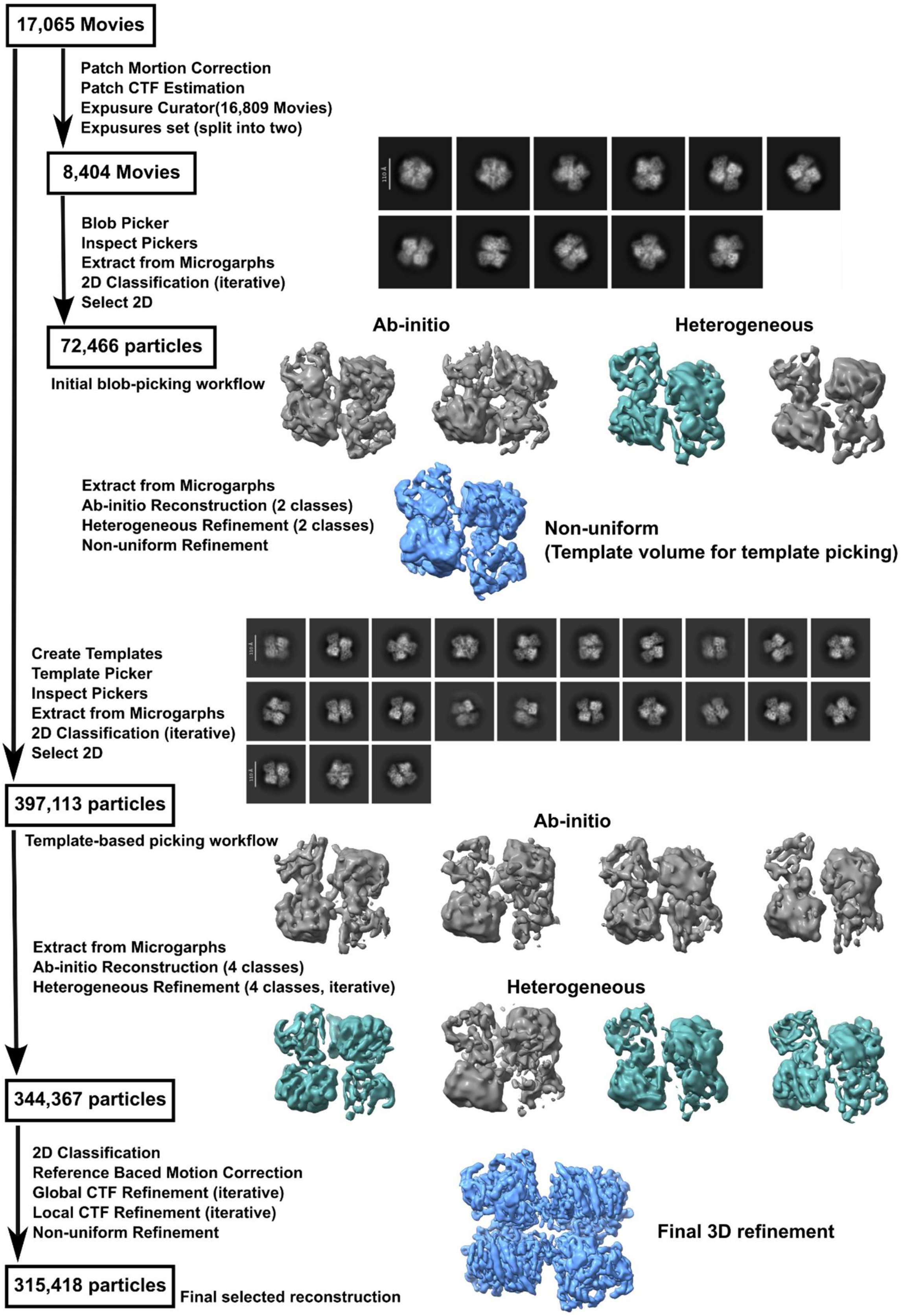

17,065 Movies
Patch Mortion Correction
Patch CTF Estimation
Expusure Curator(16,809 Movies)
Expusures set (split into two)
8,404 Movies
Blob Picker
Inspect Pickers
Extract from Micrographs
2D Classification (iterative)
Select 2D
72,466 particles
Initial blob-picking workflow
Ab-initio
Heterogeneous
Extract from Micrographs
Ab-initio Reconstruction (2 classes)
Heterogeneous Refinement (2 classes)
Non-uniform Refinement
Non-uniform
(Template volume for template picking)
Create Templates
Template Picker
Inspect Pickers
Extract from Micrographs
2D Classification (iterative)
Select 2D
397,113 particles
Template-based picking workflow
Ab-initio
Extract from Micrographs
Ab-initio Reconstruction (4 classes)
Heterogeneous Refinement (4 classes, iterative)
Heterogeneous
344,367 particles
2D Classification
Reference Baced Motion Correction
Global CTF Refinement (iterative)
Local CTF Refinement (iterative)
Non-uniform Refinement
Final 3D refinement
315,418 particles
Final selected reconstruction

**Supplementary Figure S2. CryoSPARC processing workflow for the final CYP2C9 reconstruction from the EG-grid dataset.**

Cryo-EM data collected on an epoxidized graphene grid (EG-grid) were processed in CryoSPARC. After patch motion correction, patch CTF estimation and exposure curation, an initial subset of curated micrographs was subjected to blob picking, iterative 2D classification and particle selection. Representative 2D class averages are shown. The selected particles were used for ab initio reconstruction and heterogeneous refinement to generate an initial oligomeric model, which was then used as a template for template-based particle picking on the curated dataset. After additional rounds of particle extraction, iterative 2D classification, ab initio reconstruction and heterogeneous refinement, the selected particles were further improved by reference-based motion correction, global and local CTF refinement, and non-uniform refinement. This workflow yielded the final CYP2C9 hexamer map used for structure determination. Gray maps indicate intermediate or rejected classes, cyan maps indicate selected intermediate classes, and the blue map indicates the final refined reconstruction. Scale bar in the 2D class averages, 110 Å.

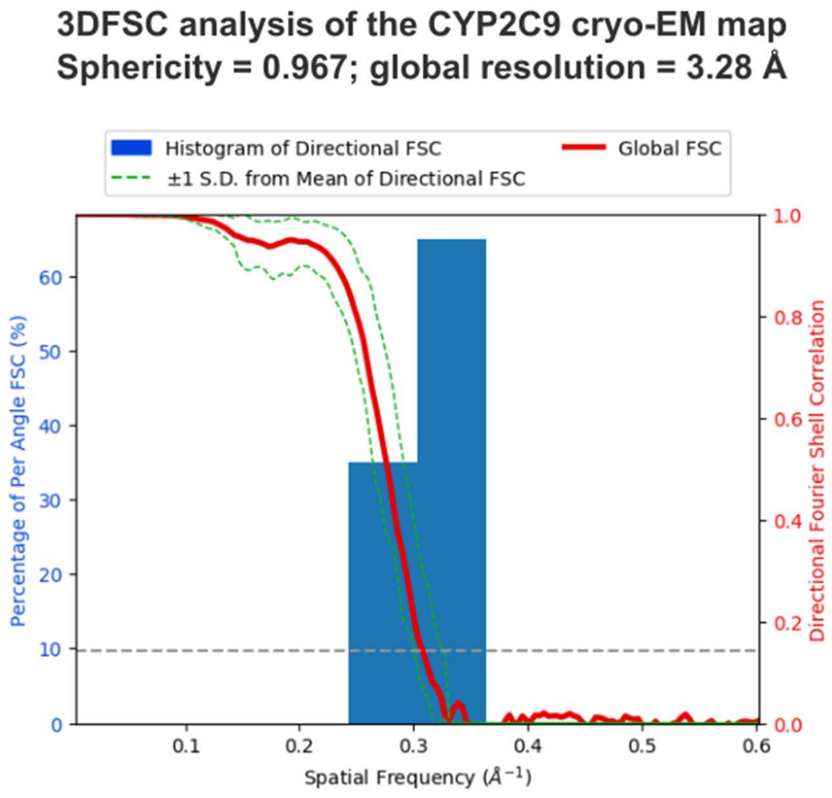


**Supplementary Figure S3. Directional FSC analysis of the CYP2C9 cryo-EM map.**

Histogram and directional FSC plot calculated using 3DFSC analysis. The analysis gave a sphericity value of 0.967 and a global resolution of 3.28 Å. The high sphericity value indicates that the reconstruction does not show severe directional anisotropy and is suitable for interpretation of the overall hexameric architecture.

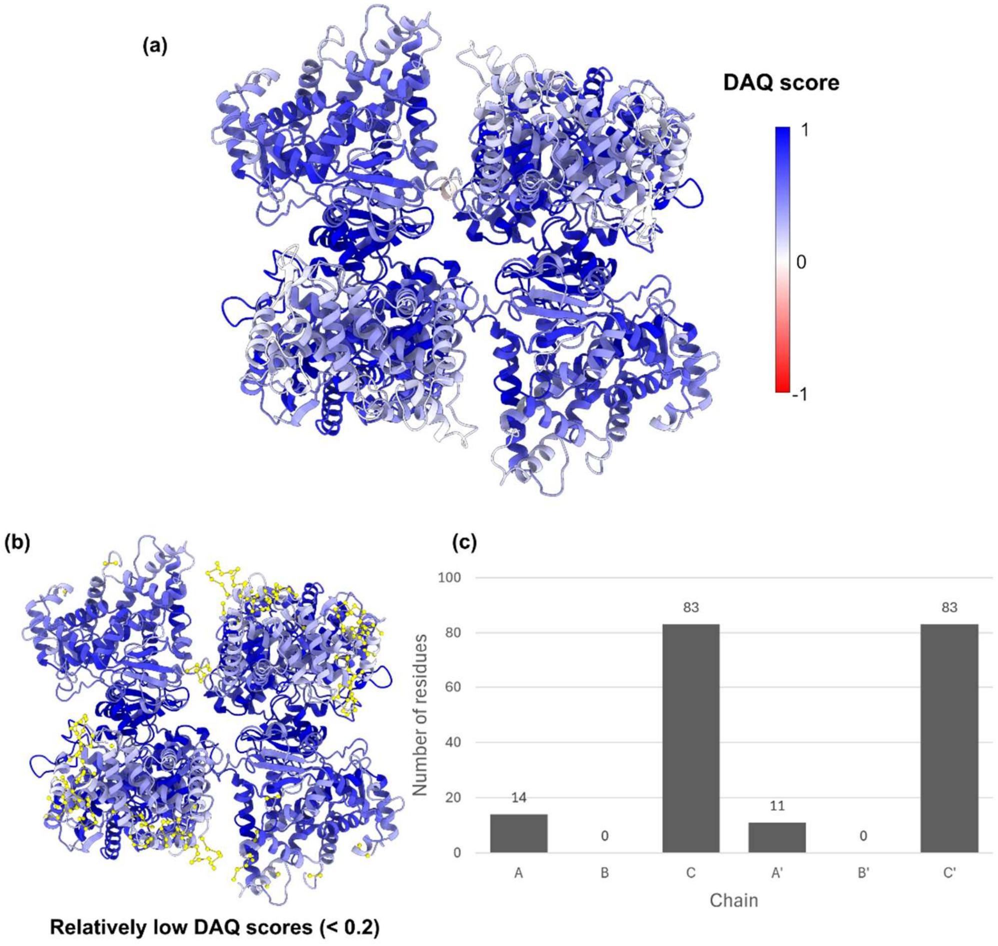


**Supplementary Figure S4. DAQ-score analysis of the CYP2C9 hexamer model.**

(a) The CYP2C9 hexamer model colored according to residue-wise DAQ score. Blue and red indicate high and low DAQ scores, respectively. No residues showed strongly negative DAQ scores (DAQ $< -0.5$). (b) Residues with relatively low DAQ scores (DAQ $< 0.2$) are highlighted in yellow. These residues were mainly located in chains C and C′. (c) Chain-wise counts of residues with DAQ scores below 0.2. Relatively low DAQ-score residues were enriched in chains C and C′, whereas no such residues were observed in chains B and B′. This distribution was consistent with the local resolution variation shown in Supplementary Fig. S5.

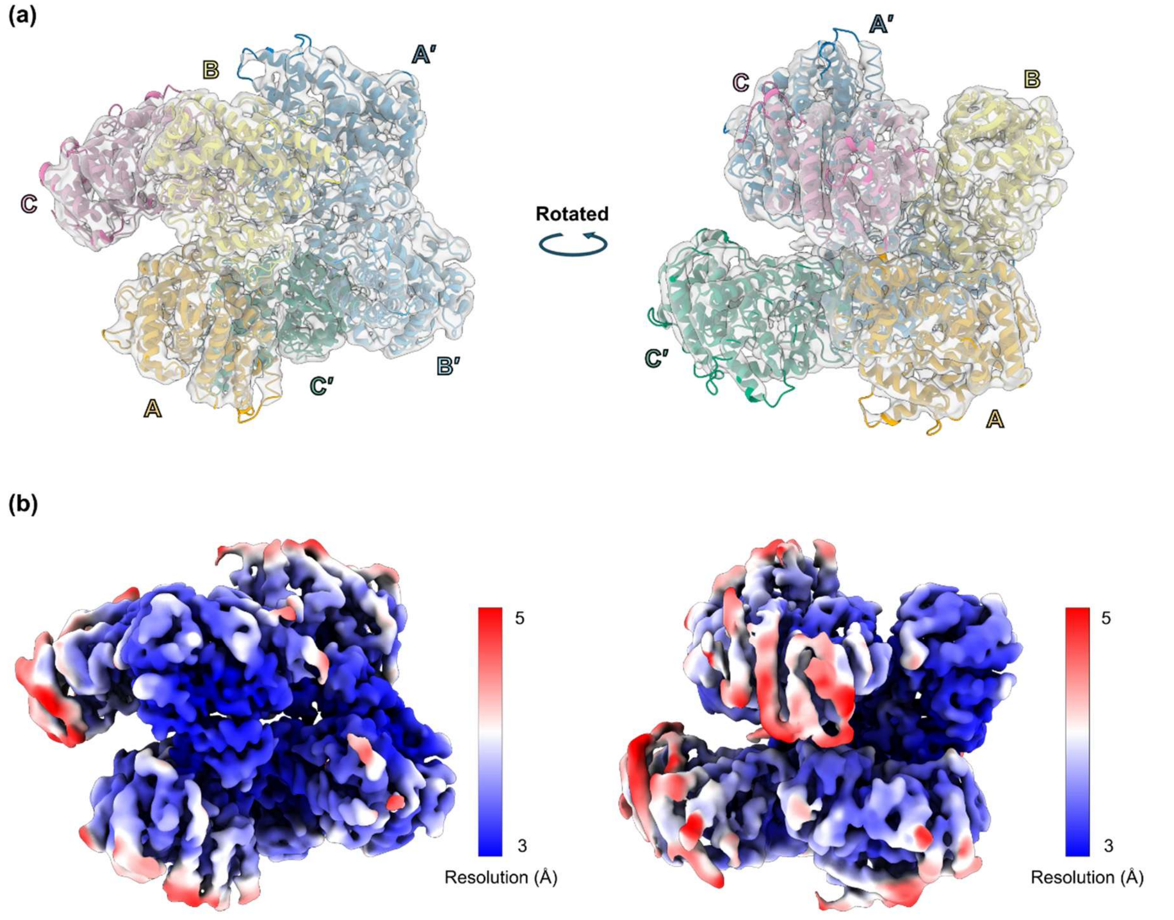


**Supplementary Figure S5. Local-resolution distribution across the CYP2C9 hexamer.**

(a) Chain assignment of the CYP2C9 hexamer shown in two orientations. Individual protomers are labeled A–C and A′–C′. (b) Local-resolution maps shown in the same orientations as in (a). Regions corresponding to chain C exhibit relatively lower local resolution than the other protomers.

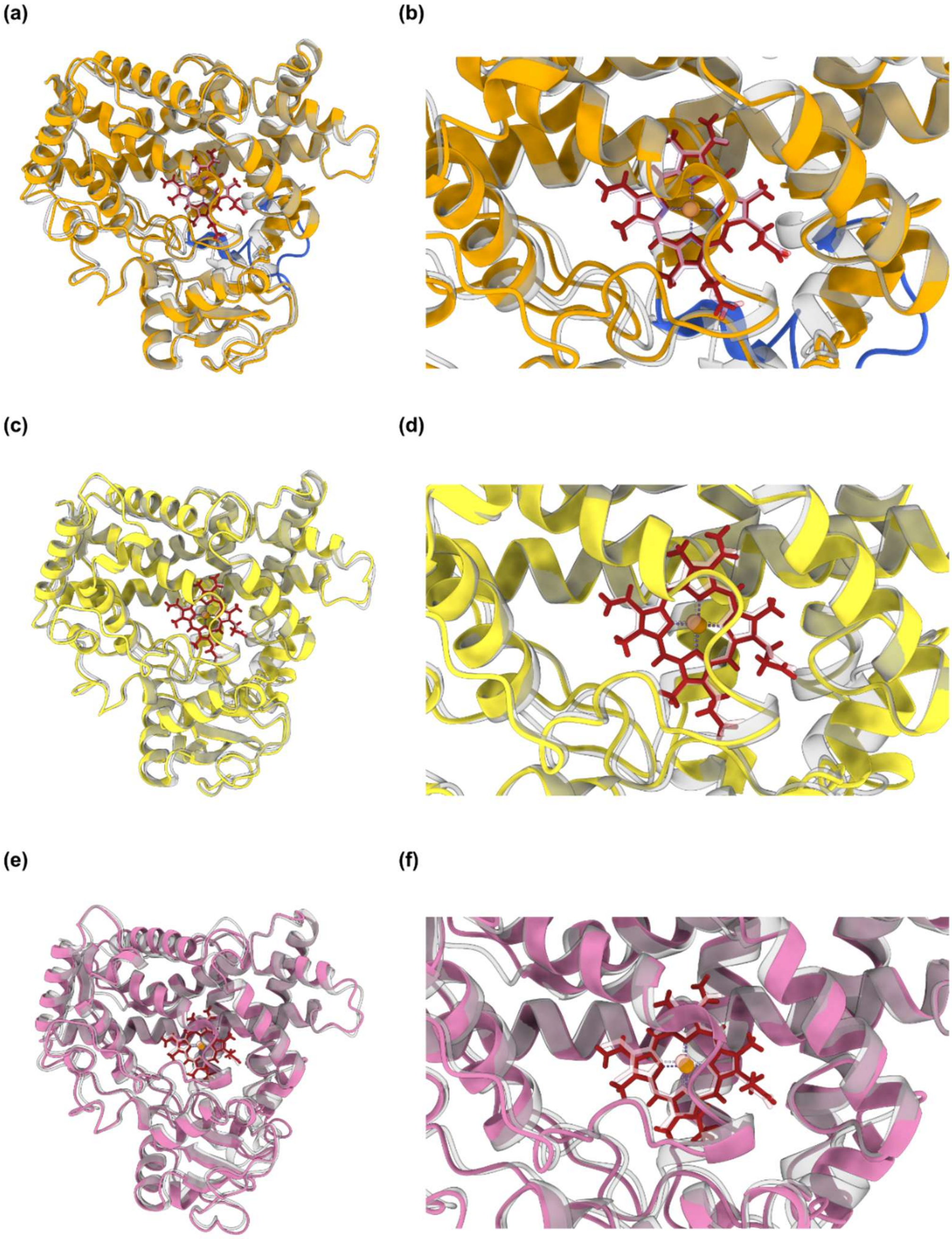


**Supplementary Figure S6. Comparison of each subunit in the cryo-EM CYP2C9 trimer structure with the crystal structure.**

Cryo-EM chains A, B and C were independently superposed onto chain A of the CYP2C9 crystal structure (PDB ID: 1OG2). Overall superpositions are shown for chain A (a), chain B (c) and chain C (e), with corresponding close-up views of the heme-binding region shown in (b), (d) and (f), respectively. The cryo-EM chains A, B and C are shown in orange, yellow and pink, respectively, and the crystal structure is shown in gray. Heme groups are shown as red sticks, with the iron atom shown as a sphere. (a) The surface-exposed region around residues 211–230 in chain A is highlighted in blue. This region shows local deviation from the crystal structure and includes residues involved in intermolecular contacts.

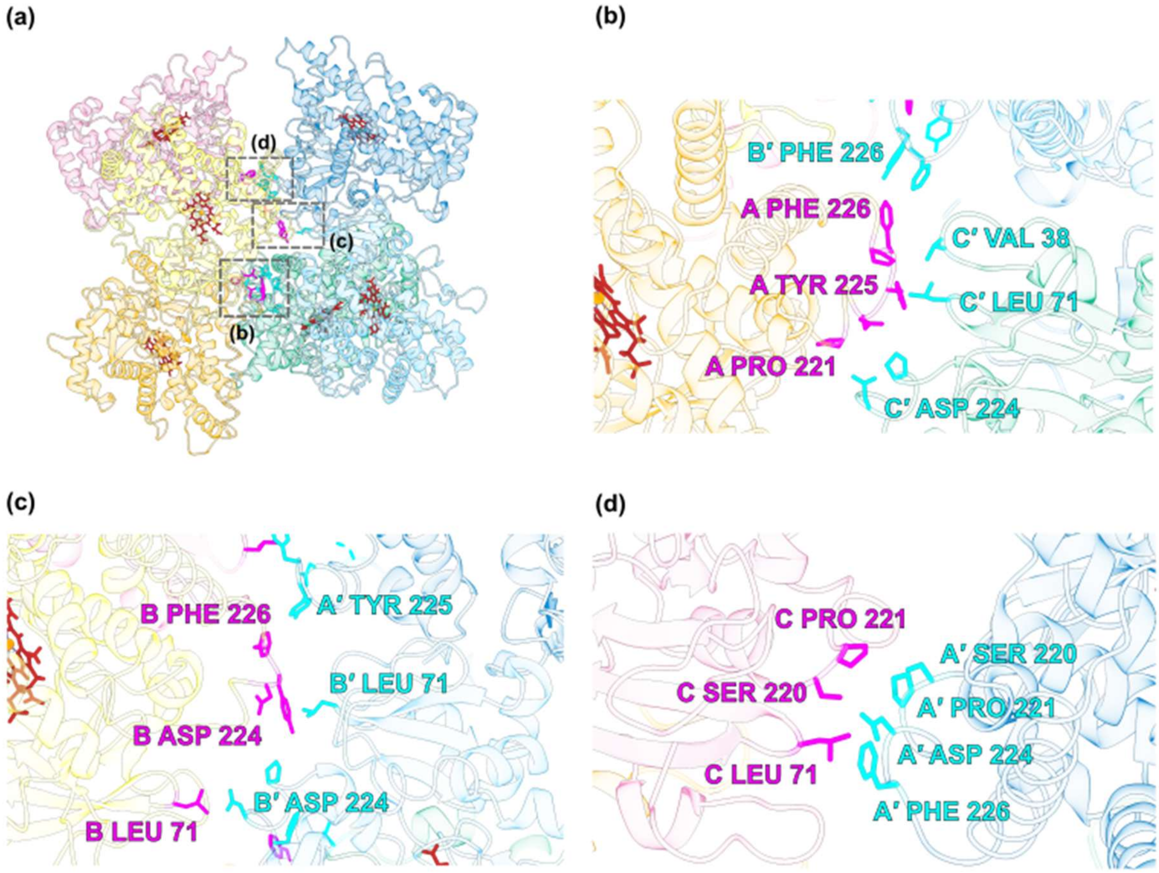


**Supplementary Figure S7. Pairwise inter-trimer contacts in the CYP2C9 hexamer.**

(a) Overview of the CYP2C9 hexamer showing the regions enlarged in panels (b)-(d). Each protomer is shown in a different color, and heme groups are shown as red sticks. Boxed regions indicate the chain A-, chain B- and chain C-centered inter-trimer contacts shown in the close-up views. (b) Chain A-centered inter-trimer contact. Chain A primarily contacts chain C′ and makes an additional limited contact with chain B′. Representative interface residues are shown as sticks and labeled. (c) Chain B-centered inter-trimer contact. Chain B primarily contacts chain B′ and makes an additional limited contact with chain A′. Representative interface residues are shown as sticks and labeled. (d) Chain C-centered inter-trimer contact. Chain C contacts chain A′, whereas no contacts with chains B′ or C′ were detected using the same distance criterion. Representative interface residues are shown as sticks and labeled. Interface residues were identified using a 4 Å distance cutoff in PyMOL. Residues from chains A–C and chains A′-C′ are highlighted in magenta and cyan, respectively.

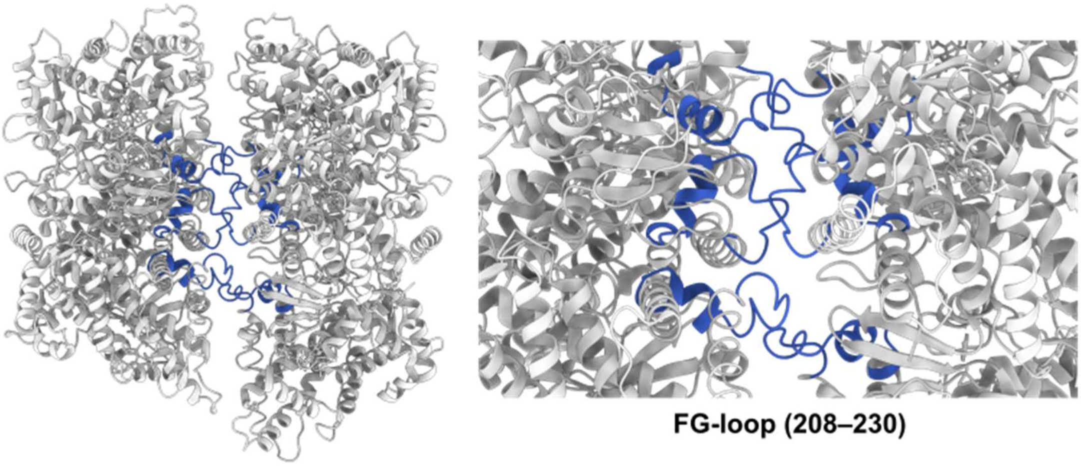


**Supplementary Figure S8. Position of the FG-loop region relative to the inter-trimer interface in the CYP2C9 hexamer.**

Overview (left) and close-up (right) views of the CYP2C9 hexamer are shown. The overall hexamer is shown in gray, and residues 208–230, corresponding to the FG-loop region, are highlighted in royal blue. The FG-loop region is located near the inter-trimer interface between the two opposing trimers.

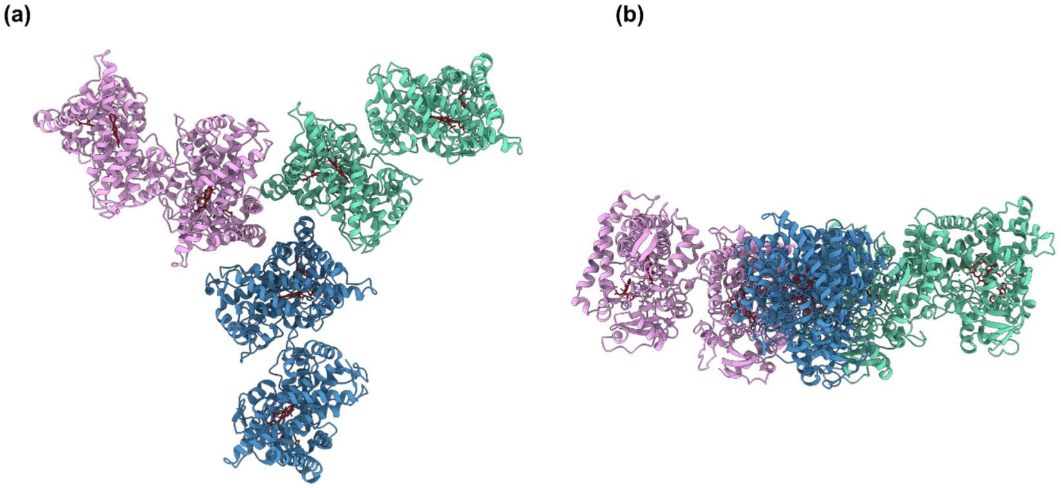


**Supplementary Figure S9. Previously proposed crystal-lattice-derived hexameric model of CYP2C9 based on PDB entry 1OG5.**

A putative CYP2C9 hexamer reconstructed from crystal contacts of PDB entry 1OG5 following the arrangement proposed by Reed and Backes (2017).

(a) View along the approximate threefold axis showing the trimer-of-dimers architecture proposed from crystal packing contacts. (b) Orthogonal view of the same assembly. The three side-to-proximal dimers are shown in different colors, and heme groups are shown in red.
